# The limitations of correlation-based inference in complex virus-microbe communities

**DOI:** 10.1101/176628

**Authors:** Ashley R. Coenen, Joshua S. Weitz

## Abstract

1

Microbes are present in high abundances in the environment and in human-associated microbiomes, often exceeding one million per milliliter. Viruses of microbes are present in even higher abundances and are important in shaping microbial populations, communities, and ecosystems. Given the relative specificity of viral infection, it is essential to identify the functional linkages between viruses and their microbial hosts, particularly given dynamic changes in virus and host abundances. Multiple approaches have been proposed to infer infection networks from time-series of *in situ* communities, among which correlation-based approaches have emerged as the *de facto* standard. In this work, we evaluate the accuracy of correlation-based inference methods using an *in silico* approach. In doing so, we compare predicted networks to actual networks to assess the self-consistency of correlation-based inference. At odds with assumptions underlying its widespread use, we find that correlation is a poor predictor of interactions in the context of viral infection and lysis of microbial hosts. The failure to predict interactions holds for methods which leverage product-moment, time-lagged, and relative-abundance based correlations. In closing, we discuss alternative inference methods, particularly model-based methods, as a means to infer interactions in complex microbial communities with viruses.

**Importance:** Inferring interactions from population time-series is an active and ongoing area of research. It is relevant across many biological systems – in particular in virus-microbe communities, but also in gene regulatory networks, neural networks, and ecological communities broadly. Correlation-based inference – using correlations to predict interactions – is widespread. However, it is well known that “correlation does not imply causation”. Despite this, many studies apply correlation-based inference methods to experimental time-series without first assessing the potential scope for accurate inference. Here, we find that several correlation-based inference methods fail to recover interactions within *in silica* virus-microbe communities, raising questions on their relevance when applied *in situ*.

## 3 Introduction

Viruses of microbes are ubiquitous and highly diverse in marine, soil, and human-associated environments. Viruses interact with their microbial hosts in many ways. For example, they can transfer genes between microbial hosts [1, 2], alter host physiology and metabolism [3, 4], and redirect the flow of organic matter in food webs through cell lysis [5, 6]. Viruses are important parts of microbial communities, and characterizing the interactions between viruses and their microbial hosts is critical for understanding microbial community structure and ecosystem function [5, 7, 8, 9].

A key step in characterizing virus-microbe interactions is determining which viruses can infect which microbes. Viruses are known to be relatively specific but not exclusive in their microbial host range. Individual viruses may infect multiple strains of an isolated microbe or they may infect across genera as part of complex virus-microbe interaction networks [10, 11]. For example, cyanophage can infect both *Prachlaracaccus* and *Synechacaccus* which are two distinct genera of marine cyanobacteria [12]. However, knowledge of viral host range remains limited because existing experimental methods for directly testing for viral infection are generally not applicable to an entire *in situ* community. Culture-based methods such as plaque assays are useful for checking for viral infection at the strain level and permit high confidence in their results, but they are not broadly applicable as many viruses and microbes are difficult or currently impossible to isolate and culture [1]. Partially culture-independent methods, such as viral tagging [13, 14] and digital PCR [15], overcome some of these hurdles but only for particular targetable viruses and microbes. Similarly, single-cell genome analysis is able to link individual viruses to microbial hosts [16, 17, 18] but for a relatively small number of cells.

Viral metagenomics offers an alternate route for probing virus-microbe interactions for entire *in situ* communities, bypassing culturing altogether [19, 20, 21]. The viral sequences obtained from metagenomes can be analyzed directly using bioinformatics-based methods to predict microbial hosts [22, 23] although such methods may only be appropriate for a subset of viruses (phages and archaeal viruses but not eukaryotic viruses) and putative hosts (prokaryotes but not eukaryotes). Alternatively, metagenomic sampling of a community *over time* can provide estimates of the changing abundances of viral and microbial populations at high time- and taxonomic-resolution. Once these high-resolution time-series are obtained, they can be used to predict virus-microbe interactions using a variety of statistical and mathematical inference methods (see reviews [24, 25, 26, 27, 28]).

Correlation and correlation-based methods are among the most widely used network inference methods for microbial communities [25]. For example, Extended Local Similarity Analysis (eLSA) is a correlation-based method which allows for both local and time-lagged correlations [29, 30, 31] and has been used to infer interaction networks in communities of marine bacteria [32, 33]; bacteria and phytoplankton [34, 35]; bacteria and viruses [36]; and bacteria, viruses, and protists [37, 38]. In addition, several correlation-based methods have been developed to address challenges associated with the compositional nature of ‘-omics’ datasets [39, 25], including Sparse Correlations for Compositional data (SparCC) [40].

Regardless of the particular details of these methods, all correlation-based inference operates on the same core assumption: that interacting populations trend together (are correlated) and that non-interacting populations do not trend together (are not correlated). Particular correlation-based methods may relax or augment this assumption. For example, with eLSA the trends may be time-lagged [29, 30, 31]; with simple rank correlations the trends may be non-parametric; and with compositional methods like SparCC the trends may occur between ratios of relative abundances [40]. In communities with only a few populations and simple interactions, population trends may indeed be indicative of ecological mechanism. In these contexts, some correlation-based methods have been shown to recapitulate microbe-microbe interactions with limited success [25]. Typically however the challenge of inferring interaction networks applies to diverse communities and complex ecological interactions. Microbial communities often have dozens, hundreds, or more distinct populations, each of which may interact with many other populations through nonlinear mechanisms such as viral lysis, as well as be influenced by fluctuating abiotic drivers. In these contexts, the relationship between correlation and ecological mechanism is poorly understood. Often correlations do not have a simple mechanistic interpretation, a well-known adage (“correlation does not imply causation”) that is often disregarded.

Despite the challenge of interpretation, correlation-based inference methods are widely used with *in situ* datasets [29, 30, 31, 32, 33, 34, 35, 36, 37, 38, 39, 25, 40]. Benchmarking inferred networks – connecting correlations to specific ecological mechanisms – is difficult. In the context of lytic infections of environmental microbes by viruses, there is (usually) no existing “gold standard” interaction network for which to validate inferred interactions. Therefore, in this work, we take an *in silica* approach to assess the accuracy of correlation-based inference. To do this, we simulate virus-microbe community dynamics with an interaction network which is prescribed *a priari* and use it to benchmark inferred networks. Several existing studies have applied similar *in silica* approaches in the case of both microbe-microbe and microbe-virus interactions and found that simple Pearson correlation [41, 39] and several correlation-based methods [25] either fail or are inconsistent in recapitulating interaction networks. Here, we provide an in-depth assessment of the potential for correlation-based inference in diverse communities of microbes and viruses. As we show, correlation-based inference fails to recapitulate virus-microbe interactions and performs worse in more diverse communities. The failure of correlation-based inference in this context raises concerns over its use in inferring microbe-parasite interactions as well as microbe-predator and microbe-microbe interactions more broadly.

## 4 Methods

### 4.1 Dynamical model of a virus-microbe community

We model the ecological dynamics of a virus-microbe community with a system of non-linear differential equations:

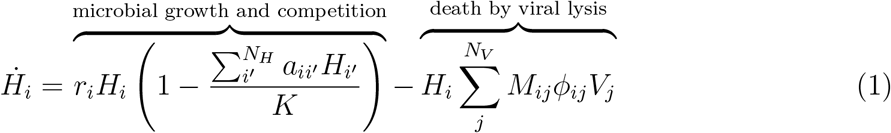

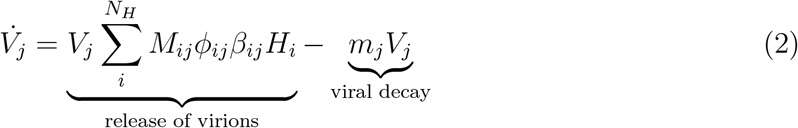

where *H_i_* and *V_j_* refer to the population density of microbial host i and virus j respectively. There are *N_H_* different microbial host populations and *N_V_* different virus populations. For our purposes, a “population” is a group of microbes or viruses with identical life history traits, that is microbes or viruses which occupy the same functional niche.

In the absence of viruses, the microbial hosts undergo logistic growth with growth rates *r_i_*. The microbial hosts have a community-wide carrying capacity *K*, and they compete with each other for resources both inter- and intra-specifically with competition strength *a_ii′_*. Each microbial host can be infected and lysed by a subset of viruses determined by the interaction terms *M_ij_*. If microbial host i can be infected by virus *j, M_ij_* = 1; otherwise *M_ij_* = 0. The collection of all the interaction terms is the interaction network represented by matrix **M** of size *N_H_* by *N*_V_. The adsorption rates *ϕ_ij_* denote how frequently microbial host *i* is infected by virus *j*.

Each virus *j*’s population grows from infecting and lysing their hosts. The rate of virus *j*’s growth is determined by its host-specific adsorption rate *ϕ_ij_* and host-specific burst size *β_ij_*, which is the number of new virions per infected host cell. The quantity 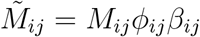 is the effective interaction strength between virus *j* and host *i*, and the collection of all the interaction strengths is the weighted interaction network 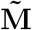. Finally, the viruses decay at rates *m_j_*.

### 4.2 Generating interaction networks and characterizing network structure

Virus-microbe interaction networks, denoted **M**, are represented as bipartite networks or matrices of size *N_H_* by *N_V_* where *N_H_* is the number of microbial host populations and *N_V_* is the number of virus populations. The element *M_ij_* is 1 if microbe population i and virus population *j* interact and 0 otherwise. In this paper, we consider only square networks (*N* = *N_H_* = *N_V_*) although the analysis is easily extended to rectangular networks. We consider three network sizes *N* = 10, 25, 50.

For each network size *N*, we generate an ensemble of networks varying in nestedness and modularity (Fig 1). We first generate the maximally nested (Fig 1A) and maximally modular (Fig 1B) networks of size *N* using the BiMat Matlab package [42]. In order to achieve maximal nestedness and modularity, the network fill *F* (fraction of interacting pairs) is fixed at *F* = 0.55 for the nested networks and *F* = 0.5 for the modular networks. For the modular networks, the number of modules is set to 2, 5, and 10 for the three network sizes respectively.

**Figure 1:**
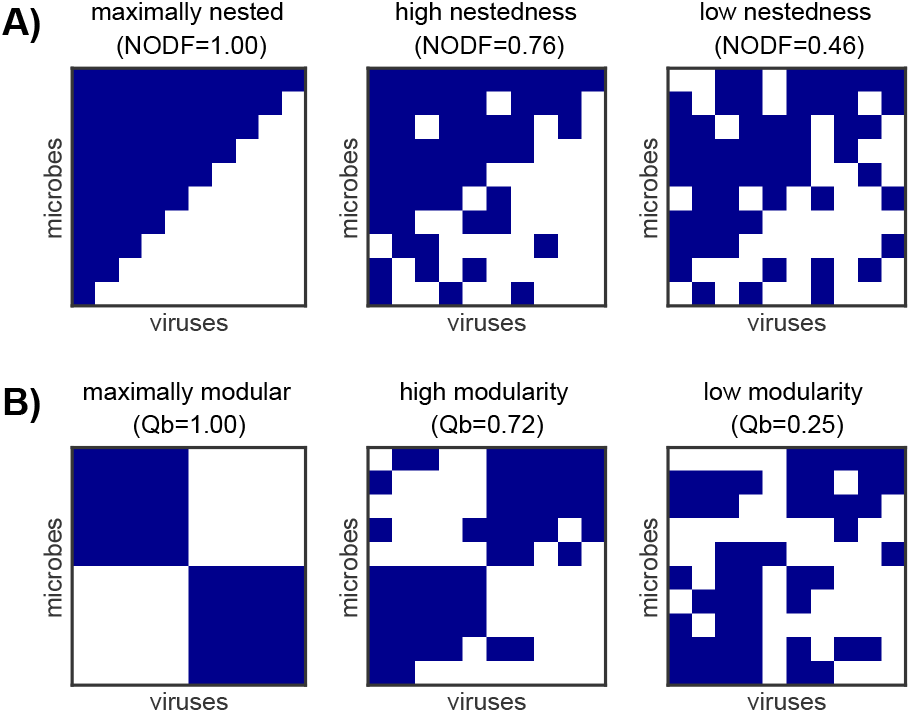
Example interaction networks characterized by A) nestedness and B) modularity. The networks shown here have size *N* =10 and fill A) *F* = 0.55 and B) *F* = 0.5. Within each network, rows represent microbe populations and columns represent virus populations, while navy squares indicate interaction (*M_ij_* = 1). Networks were generated according to §4.2. Nestedness (NODF) and modularity (*Q_b_*) were measured with the BiMat package and are arranged in their most nested or most modular forms [42].

To generate networks that vary in nestedness and modularity, we perform the following “rewiring” procedure. Beginning with the maximally nested or maximally modular network, we randomly select an interacting virus-microbe pair (*M_ij_* = 1) and a non-interacting virus-microbe pair (*M_i′j′_* = 0) and exchange their values. We do not allow exchanges that would result in an all-zero row or column, as that would isolate the microbe or virus population from the rest of the community. We continue the random selection of pairs without replacement until the desired nestedness or modularity has been achieved. To calculate nestedness and modularity, we use the default algorithms in the BiMat Matlab package. The nestedness metric used is NODF [43], and the algorithm used to calculate modularity is AdaptiveBRIM [44]. The modularity is additionally normalized according to a maximum theoretical modularity as detailed in [45].

### 4.3 Choosing life history traits for coexistence

The life history traits for a given interaction network are chosen to ensure that all microbial host and virus populations can coexist, adapted from [46].

First we sample target fixed point densities 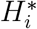 and 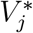 for each microbial host and virus population. In addition we sample adsorption rates *ϕ_ij_* and burst sizes *β_ij_*. All of these parameters are independently and randomly sampled from uniform distributions with biologically feasible ranges specified in Table 1. We use a fixed carrying capacity density *K* = 10^6^ cells/mL for all parameter sets.

**Table 1:** Sampling ranges for parameters in the virus-microbe dynamical model (Eqns 1 and 2).

|  | parameter | sampling range | units |
| --- | --- | --- | --- |
| $H_i^*$ | host $i$ target steady-state density | $10^3 - 10^4$ | cells/mL |
| $V_j^*$ | virus $j$ target steady-state density | $10^6 - 10^7$ | virions/mL |
| $K$ | community-wide host carrying capacity | $10^6$ | cells/mL |
| $\phi_{ij}$ | adsorption rate of virus $j$ into host $i$ | $10^{-7} - 10^{-6}$ | mL/ (virion · day) |
| $\beta_{ij}$ | burst size of virus $j$ per host $i$ | $10 - 100$ | virions/cell |
| $H_i^{0*}$ | host $i$ target steady-state density<br>in the absence of viruses | $10^3 - 10^6$ | cells/mL |
| $a_{ii'}$ | competitive effect of host $i'$ on host $i$ | $0 - 1$ | |

Next we sample microbe-microbe competition terms *a_ii′_*. We introduce an additional constraint that microbial populations should coexist in the absence of all viruses. To this end, we sample target virus-free fixed point densities 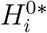 from a uniform distribution with a range specified in Table 1. After sampling, the 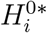 remain fixed. According to Eqn 1, coexistence in the virus-free setting is satisfied when

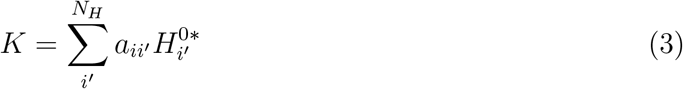

for each microbial host *i*. To start, we set all intraspecific competition to one (*a_ii_* = 1) and all interspecific competition to zero (*a_ii′_* = 0 for *i′* ≠ *i*). Then for each microbial host *i* we randomly choose an index *k* ≠ *i* and sample *a_ik_* uniformly between zero and one. If the updated sum in Eqn 3 does not exceed the carrying capacity *K*, we repeat for a new index *k*. Once the carrying capacity is exceeded, we adjust the most recent *a_ik_* so that Eqn 3 is satisfied exactly.

Finally, the viral decay rates *m_j_* and host growth rates *r_i_* are computed from the fixed point versions of Eqns 1 and 2:

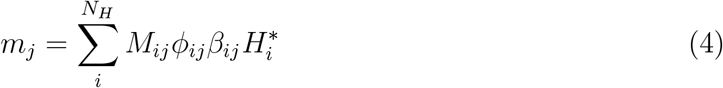

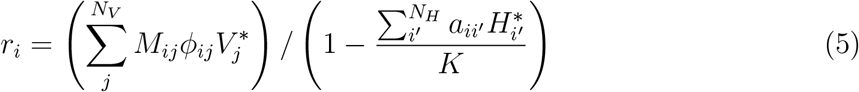

### 4.4 Simulating and sampling time-series

We use Matlab’s native ODE45 function to numerically simulate the virus-microbe dynamical model specified in §4.1 with interaction network and life history traits generated as described in §4.2 and §4.3. We use a relative error tolerance of 10^−8^. Initial conditions are chosen by perturbing the fixed point densities 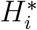 and 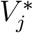 by a multiplicative factor *δ* where the sign of *δ* is chosen randomly for each microbial host and virus population. We note that *δ* can be used to tune the amount of variability in the simulated time-series (see Fig S1).

After simulating virus and microbe time-series, we sample the time-series at regularly spaced sample times (every 2 hours) for a fixed duration (200 hours, or 100 samples). Therefore, for each virus and each microbe in the community we take S samples at times *t*_1_, …, *t_S_*. We use the same sampling frequency and the same S for each inference method, except for time-delayed correlation (see §4.5).

### 4.5 Standard and time-delayed Pearson correlation networks

We assume *S* regularly spaced sample times *t*_1_, …, *t_S_* for each host type *H_i_* and each virus type *V_j_*. The samples are log-transformed, that is *h_i_*(*t_k_*) = log_10_ *H_i_*(*t_k_*) and *v_j_*(*t_k_*) = log_10_ *V_j_*(*t_k_*) for each sampled time-point *t_k_*. The standard Pearson correlation coefficient between host *i* and virus *j* is then

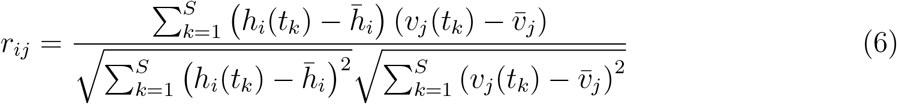

where 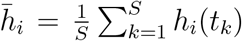 and 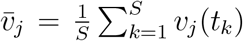 are the sample means. The correlation coefficients for all virus-host pairs are represented as a bipartite matrix **R** of size *N_H_* × *N_V_* analogous to the interaction network (see §4.2).

Time-delayed correlations are computed by sampling the virus time-series later in time. Each virus-host pair may have a unique time-delay *τ_ij_*. For example, if host *i* is sampled at times *t*_1_, …, *t_S_* then virus *j* is sampled at times *t*_1_ + *τ_ij_*, …, *t_S_* + *τ_ij_*. We keep the number of samples *S* fixed, and consequently allow virus *j* to be sampled beyond the final sample time *t_S_* of the hosts. The time-delayed Pearson correlation coefficient is

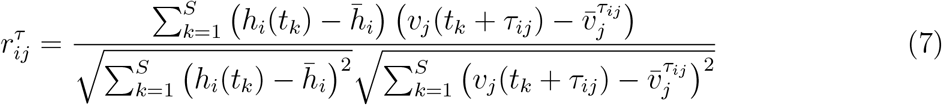

where 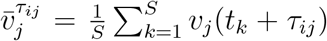 is the mean of the time-delayed virus sample. As before, the correlation coefficients for all virus-host pairs is a bipartite matrix **R**^*τ*^ of size *N_H_* × *N_V_*.

Pearson correlation coefficients, as specified above, were computed using Matlab’s native `corr` function with type=“pearson”. Alternate correlation types including Spearman and Kendall are also supported by the `corr` function and are utilized in the SI.

### 4.6 eLSA networks

Extended Local Similarity Analysis (eLSA) is a correlation-based inference method which is widely used with *in situ* time-series of complex microbial communities (*e.g*. [32, 33, 34, 35, 36, 37, 38]). eLSA attempts to detect local correlations, that is, time-series which trend together for only a portion of the sample period. In addition, eLSA allows for time-delayed correlations (as described in the previous section §4.5). To this end, a “local similarity” (LS) score is computed for each pair of time-series. The LS score is analogous to computing the Pearson correlation for all possible subsections of the two time-series, with offsets up to a pre-decided length, and keeping the maximum absolute correlation. As an example, two time-series may trend strongly during the first half of the sample period but not during the second. For such a pair of time-series, the Pearson correlation would be low, but the LS score would be high.

To compute the LS score, the two time-series are first transformed to have normal distributions (we note that such a transformation is non-stationary and thus may induce spurious correlations). The LS score is the maximal sum of the product of the entries across all possible subsections, normalized by the time-series length. If a pre-defined delay is specified, the subsections are additionally offset from one another from zero up to to the delay amount [29, 30, 31].

We applied eLSA to our simulated time-series data. We used samples of all *N_H_* host types and all *N_V_* virus types with S regularly spaced sample times *t*_1_, …, *t_S_* as input. We used the `lsa-compute.py` Python script and set parameters to specify the number of sampled points (spotNum=S), number of replicates (repNum=1), number of bootstraps (b=0), and number of permutations (x=1). All other parameters were left with their default settings including the maximum allowed time delay (delayLimit=3). The `lsa-compute.py` script computes eLSA scores between all virus-host, host-host, and virus-virus pairs. We selected only the virus-host eLSA scores and arranged them in a bipartite matrix of size *N_H_* × *N_V_* analogous to the interaction network (see §4.2). We used a custom Matlab script `write_elsa.m` to generate .csv data files in the format specified by the eLSA documentation. We used a custom bash script `elsa_compute_all.sh` to run the eLSA analysis on the ensemble of virus-microbe communities. Finally, we used a custom Matlab script `read_elsa.m` to import the results into Matlab for scoring (see §4.8).

### 4.7 SparCC networks

Sparse Correlations for Compositional data (SparCC) is a correlation-based inference method for use with compositional time-series data. This is relevant for ‘-omics’ data in which abundances are typically relative. It is well known that compositional data pose challenges for standard statistics, including Pearson and other types of correlation. Because the data sum to one, individual time-series are not independent. This biases correlations to be negative regardless the trend between the underlying absolute abundances. SparCC estimates the Pearson correlation between two time-series while taking into account these compositional dependencies. In particular, SparCC computes the variance of the log-transformed ratio of two time-series, and compares this quantity to the variances of the individual log-transformed time-series. SparCC assumes sparsity in the correlation matrix but is robust to violations of this assumption [40].

We applied SparCC to our simulated time-series data as a means to evaluate correlation-based inference in a scenario in which underlying viral and microbial densities can be measured only relatively. Given samples at *S* regularly spaced sample times *t*_1_, …, *t_S_*, we first normalized the *N_H_* host types and *N_V_* virus types at each sample time *t_k_* by

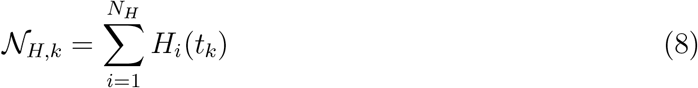

for the hosts and by

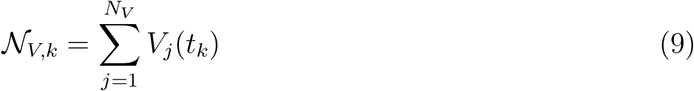

for the viruses. We used the normalized *N_H_* host and *N_V_* virus samples as input for the SparCC computation using the `SparCC.py` script. All parameters were left with their default settings. We used a custom Matlab script `write_sparcc.m` to generate .csv data files in the format specified by the SparCC documentation. We used a custom bash script `sparcc_compute_all.sh` to run the SparCC analysis on the ensemble of virus-microbe communities. Finally, we used a custom Matlab script `read_sparcc.m` to import the results into Matlab for scoring (see §4.8).

### 4.8 Scoring correlation network accuracy

To evaluate how well the Pearson correlation, eLSA, or SparCC (collectively referred to as “correlation”) network **R** recapitulates the original interaction network 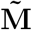, we compute the receiving operator curve (ROC). First, we binarize the interaction network 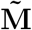 so that it is a boolean network **M** of zeros (non-interactions) and ones (interactions). Then we choose a threshold of interaction *c* between the minimum and maximum attainable values of the correlation network **R**; for Pearson correlation these are −1 and +1. Correlations in **R** that are greater than or equal to c are categorized as interactions (ones), while those that are less are non-interactions (zeros). The true positive (TP) count is the number of interactions in **M** correctly predicted by the thresholded correlation network **R_c_**. The false positive (FP) count is the number of non-interactions in **M** incorrectly predicted by **R_c_**. The TP and FP counts are normalized by the number of interactions and non-interactions in **M** to obtain the true positive rate (TPR) and false positive rate (FPR). TPR and FPR are computed for all thresholds c to obtain the receiver operator curve (ROC).

The overall “score” of the correlation network **R** is the area under the curve (AUC). A perfect prediction results in AUC=1, since for some threshold TPR=1 and FPR=0. Random predictions result in AUC=1/2, since TPR=FPR across all possible thresholds. AUC values which are less than 1/2 indicate a misclassification of “interaction”, that is, categorizing interactions and non-interactions in the opposite way would have resulted in a better prediction of 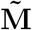.

## 5 Results

### 5.1 Standard Pearson correlation

We calculated the standard Pearson correlation networks for an ensemble *in silico* communities that varied in network size and network structure. For each network size *N* = 10, 25, 50, we generated 20 unique interaction networks. 10 of the networks were generated so that they were distributed along a range of nestedness values, and the other 10 were generated so that they were distributed along a range of modularity values (see §4.2). For each interaction network, a single set of life history traits were generated to ensure coexistence using biologically feasible ranges according to §4.3. The mechanistic model for the community dynamics is described in §4.1. Time-series were simulated according to §4.4 with *δ* = 0.3, that is, the initial conditions were the fixed point values perturbed by 30% (for additional values of *δ* see Fig S5 in the SI). For *δ* = 0.3, the mean coefficient of variation was 12% for host time-series and 4% for virus time-series (see Fig S1 in the SI). The time-series were sampled during the transient dynamics to represent *in situ* communities which are likely perturbed from equilibrium due to changing environmental conditions and intrinsic feedback. We sampled the time-series every 2 hours for 200 hours, that is, we took 100 samples (for additional sample frequencies see Fig S7 in the SI).

For each *in silico* community, we calculated the standard Pearson correlation network as described in §4.5. Two example *in silico* communities of size N =10 are shown in Fig 2 with their simulated time-series, log-transformed samples, and resulting correlation networks. The correlation networks were scored against the original interaction networks by computing AUC as described in §4.8. The procedure for computing AUC is shown in Fig 3 for the two example *in silica* communities.

**Figure 2:**
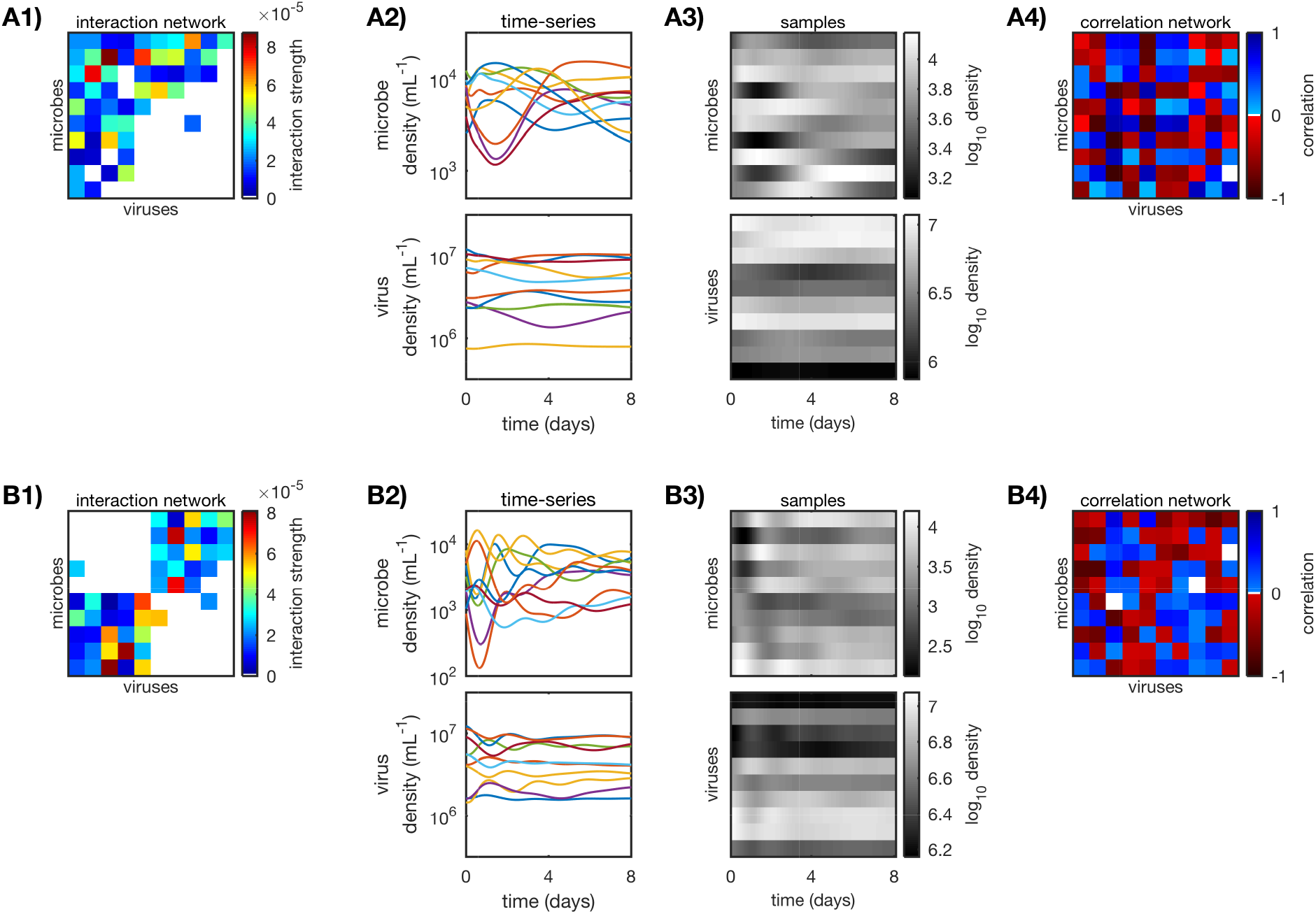
Calculating standard Pearson correlation networks for two *in silico* A) nested and B) modular communities (*N* = 10). A1-B1) Original weighted interaction networks, generated as described in §4.2 and §4.3. A2-B2) Simulated time-series of the virus-microbe dynamical system as described in §4.4 (*δ* = 0.3). A3-B3) Log-transformed samples, sampled every 2 hours for 200 hours from the simulated time-series. A4-B4) Pearson correlation networks, calculated from log-transformed samples as described in §4.5.

**Figure 3:**
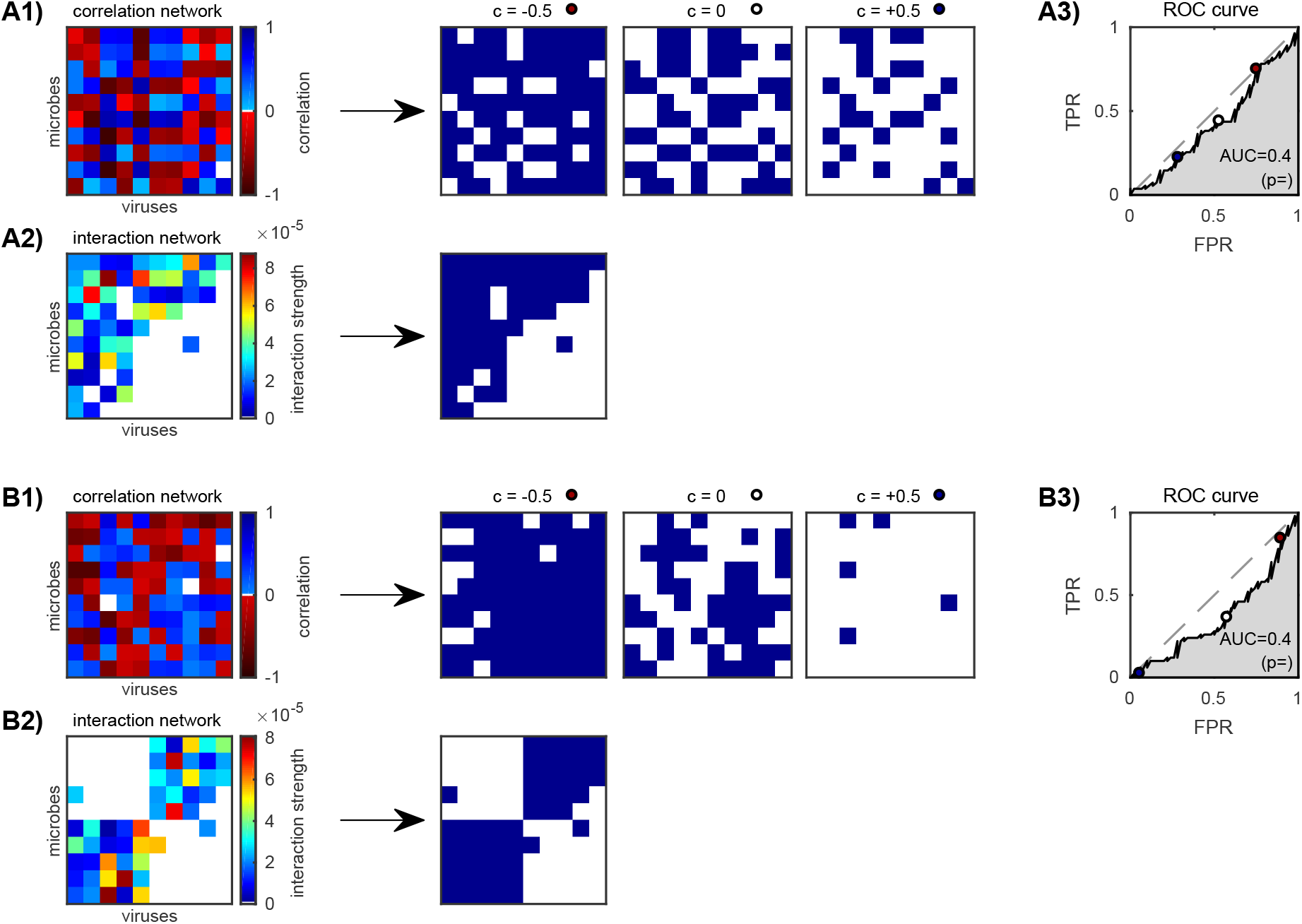
Scoring correlation network accuracy of the two *in silico* A) nested and B) modular communities (*N* = 10, see Fig 2) as described in §4.8. A1-B1) Correlation networks are binarized according to thresholds *c* between −1 and +1, three of which are shown here (*c* = −0.5,0, and 0.5). A2-B2) Original interaction networks are also binarized. A3-B3) True positive rate (TPR) versus false positive rate (FPR) of the binarized correlation networks for each threshold *c*. Three example thresholds (*c* = −0.5,0, and 0.5) are marked (red, white, and blue circles). The “non-discrimination” line (grey dashed) is where TPR = FPR. The AUC or area under the ROC curve is a measure of relative TPR to FPR over all thresholds; AUC = 1 is a perfect result. Distributions for the reported p-values are shown in the SI.

AUC values for all *in silica* communities are shown in Fig 4. Across varying network size and network structure, AUC is approximately 1/2 implying that standard Pearson correlation networks lack predictive power. Similar results were found when varying the initial condition perturbation *δ* (Fig S4) and the sampling frequency (Fig S7). There are some cases for the smaller networks (*N* = 10) where AUC does deviate from 1/2 although these deviations are small (≈ ±10%). Interestingly these deviations tend to be negative indicating a misclassification of the interaction condition, that is, negative correlations are slightly better predictors of interaction than positive correlations. Overall however, the deviations disappear for larger networks (*N* = 50) implying that they are exceptions rather than the norm. We completed identical analyses for additional correlation metrics in particular Spearman correlation and Kendall correlation (see Fig S2 in SI). We found similar results reinforcing our conclusion that simple correlations between time-series are poor predictors of the underlying interaction network.

**Figure 4:**
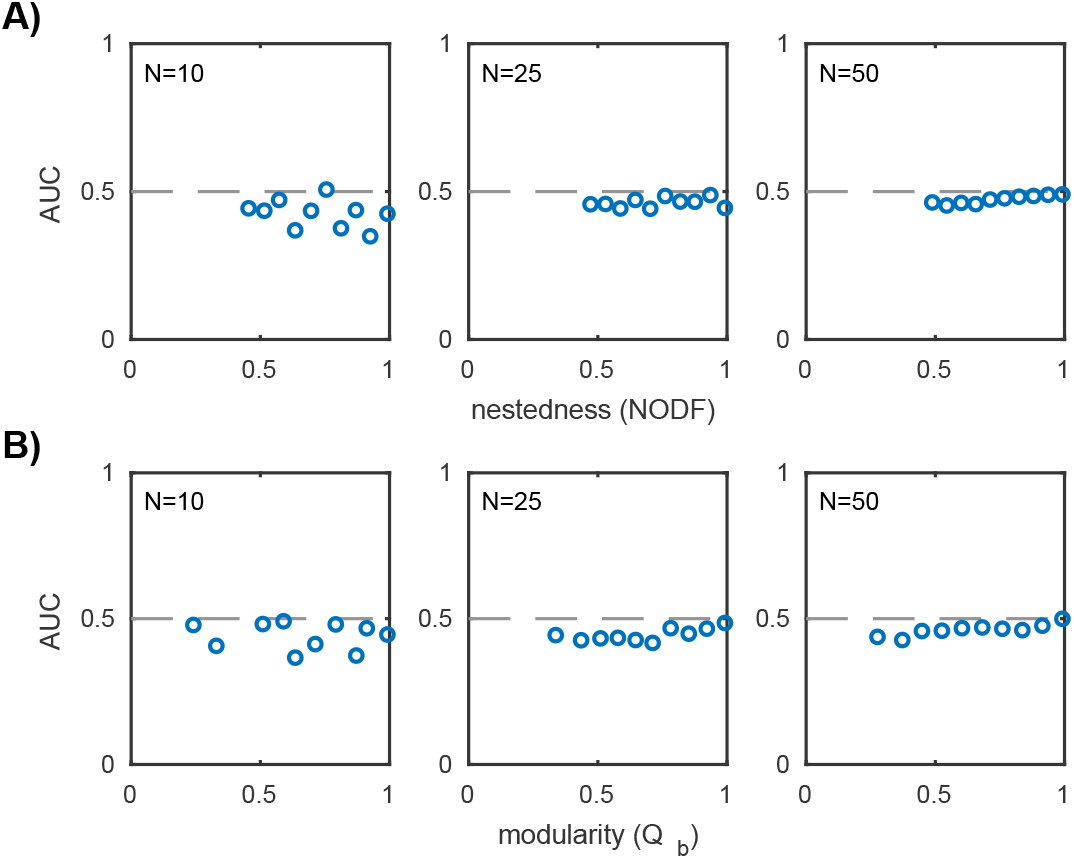
AUC values for standard Pearson correlation for the ensemble of A) nested and B) modular communities over three network sizes *N* = 10, 25, 50 (20 communities for each network size). AUC is computed as described in §4.8. Each plotted point corresponds to a unique *in silico* community. Dashed line marks AUC=1/2 and implies the predicted network did no better than random guessing.

### 5.2 Time-delayed Pearson correlation

Given the results of the previous section §5.1 – that standard correlations do not recapitulate interactions – we computed time-delayed correlation networks for the same ensemble of *in silica* communities. The addition of time-delays to standard correlation approaches is motivated by a large body of theoretical work on predator-prey dynamics, where both predator and prey populations oscillate but with a phase delay between them [47]. Similar results hold for the phase delay in simple phage-bacteria dynamics [48]. Time-delayed corre- lations are the basis of several existing correlation-based inference methods including eLSA [29, 30, 31].

For this analysis, we used the same ensemble of *in silico* communities (network sizes *N* = 10, 25, 50 of varying nestedness and modularity), simulated time-series (*δ* = 0.3; see Fig S5 in SI), and sample frequency (2 hours; see Fig S8 in SI) as before (see §5.1 for time-series). We calculated the time-delayed Pearson correlation networks as described in §4.5, where for each virus-host pair the virus time-series is sampled later in time by some delay amount *τ_ij_* relative to the host time-series (for Spearman and Kendall correlation, see Fig S3 in SI). Each delay is chosen such that the absolute value of the correlation for the virus-host pair is maximized. Since the optimal time-delay is not known in advance, delays between 0 < *τ_ij_* < *t_S_*/2, (0 hours and *t_S_*/2 = 100 hours) were considered. The number of samples used to compute each correlation coefficient was kept fixed at *S* =100 (sample duration 200 hours). Time-delayed Pearson correlation networks for the two example *in silica* communities of size *N* =10 are shown in Fig 5A-B. AUC was computed as described in §4.8.

**Figure 5:**
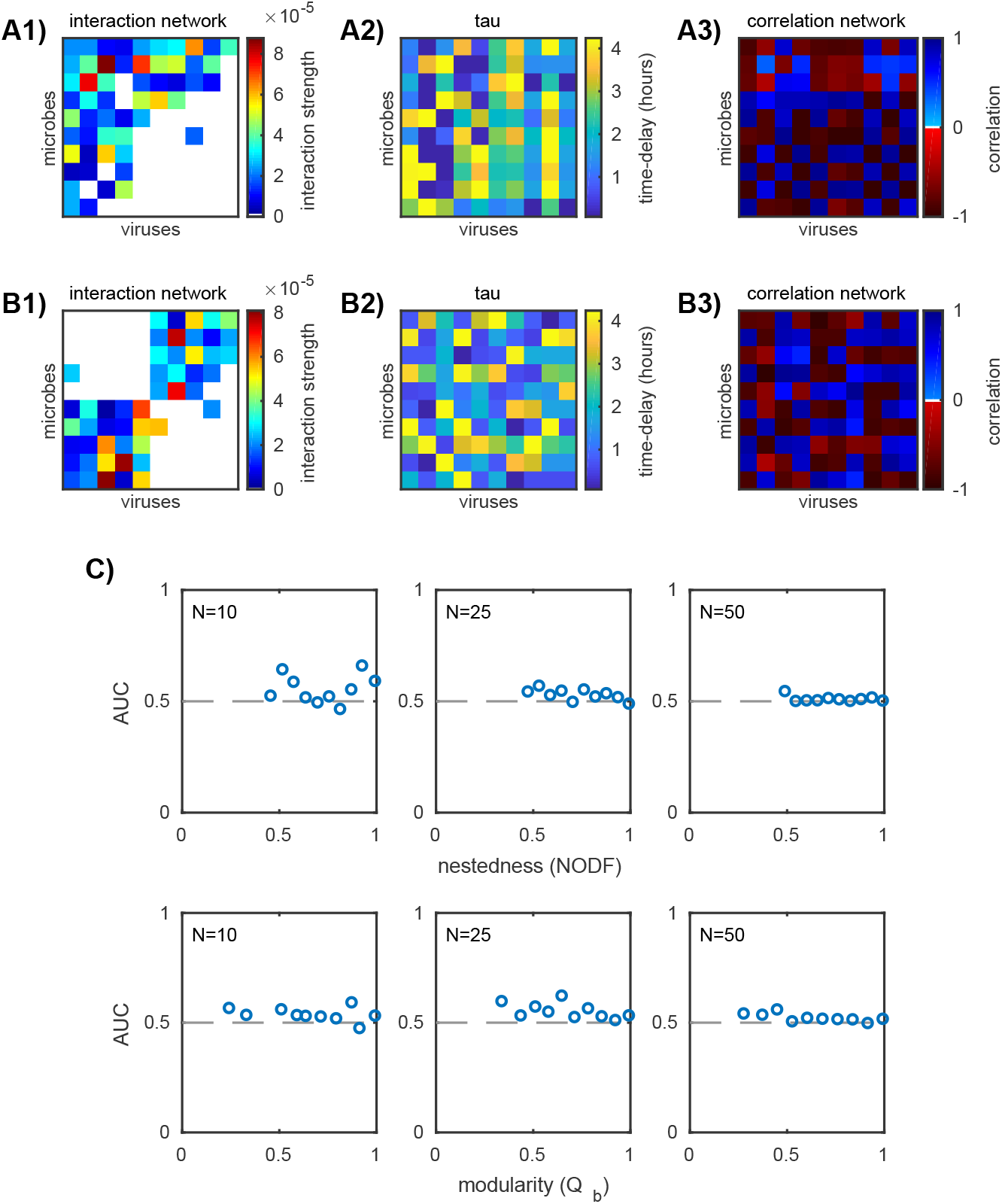
Performance of time-delayed Pearson correlation. A1-B1) Two example *in silico* interaction networks (*N* = 10). A2-B2) Time-delays *τ_ij_* for each virus-host pair, chosen so that the absolute value of the correlation is maximized. A3-B3) Time-delayed Pearson correlation networks calculated as described in §4.5. C) AUC values for the ensemble of nested (top row) and modular (bottom row) communities over three network sizes *N* = 10, 25, 50 (20 communities for each network size). Each plotted point corresponds to a unique *in silico* community. Dashed line marks AUC=1/2 and implies the predicted network did no better than random guessing.

AUC values for all *in silica* communities are shown in Fig 5C. For the small networks (*N* = 10) there are a few particular networks which have AUC scores greater than 1/2. For the remaining small networks and the large networks (*N* = 25, 50), AUC ≈ 1/2 implying time-delayed Pearson correlation lacks predictive power for these networks. Similar results were found for alternate correlation metrics (Spearman and Kendall; Fig S3), initial condition perturbations *δ* (Fig S5), and sampling frequencies (Fig S8). Because AUC deviates from 1/2 for only a few small networks and disappears for large networks, it should be considered an exceptional result rather than the norm for time-delayed Pearson correlation.

### 5.3 Correlation-based methods eLSA and SparCC

We performed a similar *in silico* analysis using eLSA [29, 30, 31] and SparCC [40], two established correlation-based inference methods which are widely used with *in situ* time-series data. We used the same ensemble of *in silico* communities as before (network sizes *N* = 10, 25, 50 of varying nestedness and modularity), along with the simulated time-series (*δ* = 0.3; see Fig S6), sample frequency (2 hours; see Fig S9) and sample duration (200 hours). We implemented eLSA and SparCC as described in §4.6 and §4.7 respectively. eLSA and SparCC predicted networks for the two example *in silico* communities of size *N* =10 are shown in Fig 6A-B. AUC was computed as before and as described in §4.8.

**Figure 6:**
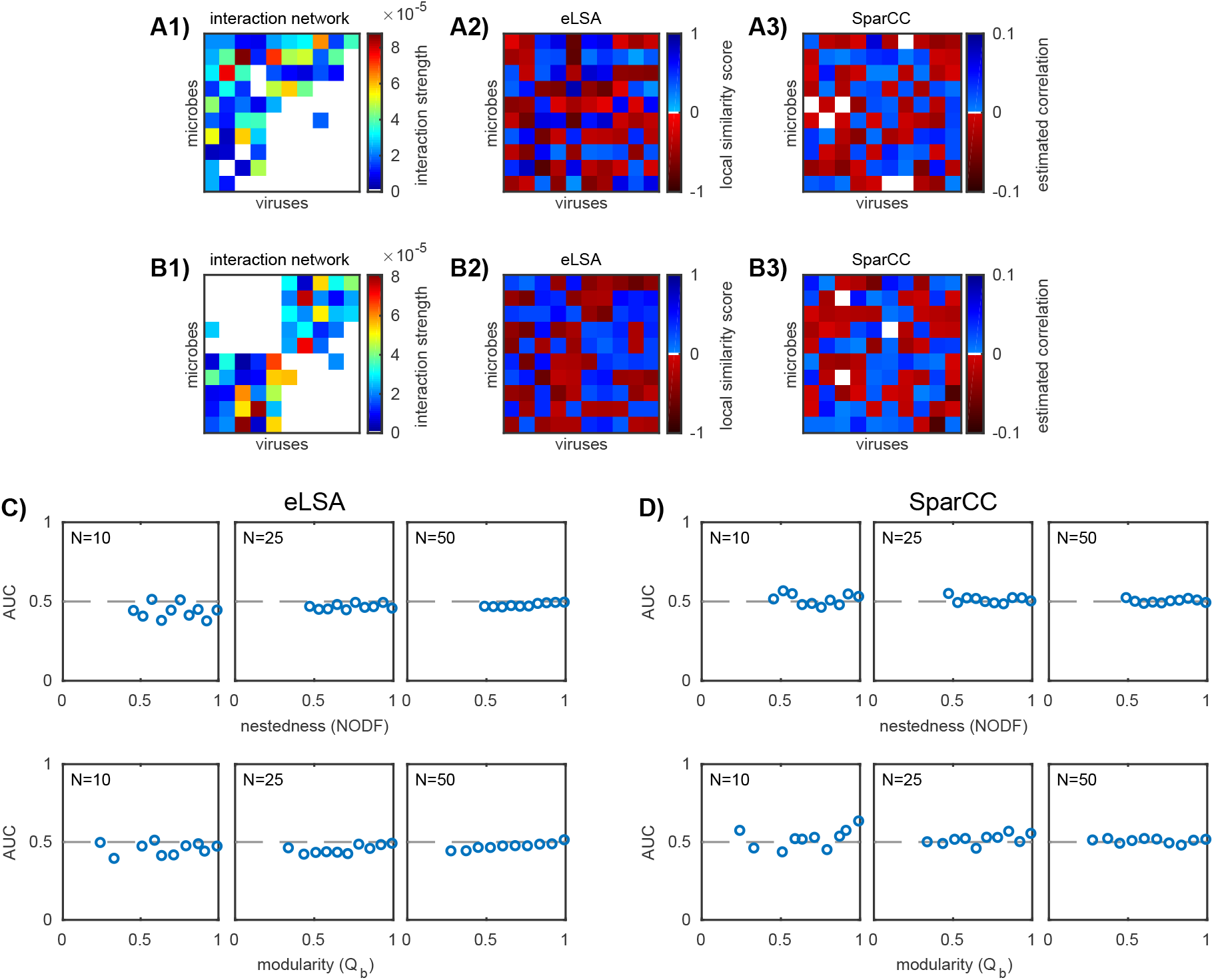
Performance of correlation-based inference methods eLSA and SparCC. A1-B1) Two example *in silico* interaction networks (*N* = 10). A2-B2) eLSA predicted network computed as described in §4.6. A3-B3) SparCC predicted network computed as described in §4.7 (color bar adjusted for visibility). C-D) AUC values for the ensemble of nested (top row) and modular (bottom row) communities over three network sizes *N* = 10, 25, 50 (20 communities for each network size). Each plotted point corresponds to a unique *in silico* community. Dashed line marks AUC=1/2 and implies the predicted network did no better than random guessing.

AUC values for all *in silica* communities are shown in Fig 6C. We see the same trends as with standard correlation and time-delayed correlation (see Figs 4 and 5). Similar results hold for varying values of the initial condition perturbation *δ* (Fig S6) and sampling frequency (Fig S9). For small networks (*N* = 10), there are a few AUC scores which deviate weakly from 1/2 (≈ ±10%). Interestingly, AUC scores for eLSA tend to be negative, implying a misclassification of interaction. AUC converges to 1/2 as network size increases (*N* = 25, 50) indicating that the AUC scores for small networks may themselves be spurious.

## 6 Discussion

Using *in silico* virus-microbe community dynamics, we calculated correlation networks among viral and microbial population time-series samples. We tested the accuracy of several different types of correlation and time-delayed correlation (Pearson, Spearman, and Kendall) and existing correlation-based inference methods (eLSA and SparCC). The correlation networks for all of these implementations failed to effectively predict the original interaction networks, as quantified by the AUC score. Failure persisted across variation in network structure, network size, degree of initial condition perturbation (*i.e*. scaling the variability of dynamics), and sampling frequency. We therefore conclude these correlation-based inference methods do not meaningfully predict interactions given this mechanistic model of virus-microbe community dynamics.

Earlier, we stated the core assumption of correlation-based inference: that interacting populations are correlated and that non-interacting populations are not correlated. While this core assumption may sometimes hold in small microbe-only communities with simple interaction mechanisms [25], we find it does not necessarily hold in more complex virus-microbe communities. (Each inference method also faces challenges unique to its formulation: eLSA in particular uses a non-stationary data transformation which may induce additional spurious correlations.) We considered communities with microbes and viruses that interacted through a nonlinear mechanism (infection and lysis) across a spectrum of network sizes and network structure. We found that correlation-based inference performed poorly given variation in these network properties, but that there was greater variation in performance for small networks. Because this variation is relatively small and disappears for larger networks, successful predictions for small networks may themselves be spurious. Namely, for a small network (*e.g. N* < 10), there is a greater probability of randomly guessing the interactions correctly because the space of possible networks is smaller.

Our results raise concerns about the use of correlation-based methods on *in situ* datasets, since a typical community under consideration will have dozens or more interacting strains and therefore will not be in the low diversity microbe-only regime explored by [25]. Additional challenges such as external environmental drivers, measurement noise, and system stochasticity must also be carefully considered before applying correlation-based methods to *in situ* datasets. Although the degree of variability of dynamics had no effect on inference quality here, it may also be an important consideration for both experimental design and choice of inference method. For example, the model-based inference method developed by [49] performs better when dynamics are highly variable. On the other hand, co-occurrence based inference methods, which require samples across space instead of time, may enable inference across different baseline environmental conditions even if the dynamics within a given environment are relatively stable.

In light of the poor performance of correlation-based methods, we advocate for increased studies of model-based inference. Model-based inference methods operate by first assuming an underlying dynamical model for the community (such as the one used in this manuscript, Eqns 1 and 2). The dynamical model is then used to formulate an objective function for an optimization or regression problem, where the solution is the interaction network which best describes the sampled community time-series (for example, see [41, 39, 50, 49, 51, 52]). Unlike correlation-based methods which assume that similar trends in population indicate interaction, model-based inference has the potential to be tailored to complex communities and environments while leveraging existing knowledge about ecological mechanisms. Given favorable results of *in silico* benchmarking of model-based inference methods [41, 39, 50, 49, 51, 52], it will be important to investigate the efficacy of model-based inference methods for complex microbial and viral communities in practice.

## 7 Acknowledgments

We are grateful to Ben Bolduc, Stephen Beckett, and five anonymous reviewers for helpful comments and feedback. We thank both Yu-Hui Lin and David Demory for reviewing the code used in the analysis. This work was supported by the Simons Foundation (SCOPE award ID 329108, J.S.W.).

## 8 Availability of data and materials

Analysis was primarily performed in Matlab. All Matlab scripts, Matlab data files (also available as .csv files), and custom bash scripts for implementing eLSA and SparCC are publicly available on GitHub (https://github.com/WeitzGroup/correlation_based_inference) and archived on Zenodo (DOI 10.5281/zenodo.844918). The BiMat Matlab package [42] used for characterizing bipartite networks is available on GitHub (https://github.com/cesar7f/BiMat). The eLSA Python package [29, 30, 31] is available on Bit-bucket (https://bitbucket.org/charade/elsa/wiki/Home). The SparCC Python package [40] is available on Bitbucket (https://bitbucket.org/yonatanf/sparcc).

